# Parkinson’s Disease Genetic Risk Evaluation in Microglia Highlights Autophagy and Lysosomal Genes

**DOI:** 10.1101/2020.08.17.254276

**Authors:** Alix Booms, Steven E. Pierce, Gerhard A. Coetzee

**Affiliations:** Center for Neurodegenerative Science, Van Andel Institute, Grand Rapids, MI, 49503, United States of America

## Abstract

Genome-wide association studies (GWAS) have uncovered thousands of single nucleotide polymorphisms (SNPs) that are associated with Parkinson’s disease (PD) risk. The functions of most of these SNPs, including the cell type they influence, and how they affect PD etiology remain largely unknown. To identify functional SNPs, we aligned PD risk SNPs within active regulatory regions of DNA in microglia, a cell type implicated in PD development. Out of 6,749 ‘SNPs of interest’ from the most recent PD GWAS metanalysis, 73 were located in open regulatory chromatin as determined by both ATAC-seq and H3K27ac ChIP-seq. We highlight a subset of SNPs that are favorable candidates for further mechanistic studies. These SNPs are located in regulatory DNA at the *SLC50A1, SNCA, BAG3, FBXL19, SETD1A*, and *NUCKS1* loci. A network analysis of the genes with risk SNPs in their promoters, implicated substance transport, involving autophagy and lysosomal genes. Our study provides a more focused set of risk SNPs and their associated risk genes as candidates for further follow-up studies, which will help identify mechanisms in microglia that increase the risk for PD.

## Introduction

Genetic studies suggest that a significant portion of Parkinson’s disease (PD) risk is heritable [1, 2]. Unlike some rare disorders caused by high penetrance mutations in a small number of genes, PD is linked to many low penetrance variants with more modest influence on disease risk. The latest genome-wide association study (GWAS) of PD has identified over 90 independent SNPs that are correlated with increased risk for the disease [2]. However, these 90 SNPs tag larger blocks of chromatin that contain many other SNPs in linkage disequilibrium. In other words, the independent SNPs can be considered surrogates for hundreds of additional SNPs that are co-inherited and therefore carry the same statistical association for risk. In total, there are thus tens of thousands of SNPs that are associated with increased risk for PD, but not all of them are causal for the disease or even functional in relevant cellular and developmental contexts. Moreover, the majority of these SNPs are located in non-coding DNA [3]. Thus, the first challenges are to 1) determine <u>*which*</u> SNPs are functional and 2) dissect the <u>*mechanisms*</u> by which each allele of a risk SNP leads to biological differences.

Multiple strategies are used to define or narrow down which SNPs are functional. In this study, we considered a SNP provisionally functional if it was located in active regulatory DNA and predicted to disrupt one or more transcription factor binding motifs. These SNPs presumably affect the expression of one or more genes through allele-specific binding of transcription factors. A confirmation of this general mechanism was demonstrated by Soldner et al. [4]. The authors found that two transcription factors, EMX2 and NKX6-1, had a lower binding preference for the risk allele of a SNP located within an enhancer at the SNCA gene. In turn, this led to a slight increase in SNCA expression. For nearly all of the thousands of other PD risk SNPs, similar mechanisms have yet to be evaluated.

A SNP may be functional in one cell type but not another, depending on the cell’s regulatory landscape. It is therefore essential to assess genetic risk in a cell-type-specific context. For PD, there are multiple cell types, other than dopaminergic neurons, that are likely involved in disease development and progression. Cell types like astrocytes, oligodendrocytes, and microglia express PD-associated lysosomal gene sets at higher levels than neuronal cell types [5]. Immune cells of the myeloid lineage, including CNS resident microglia, also highly express genes at PD associated loci [6-8], suggesting that glia and immune cell types are candidates in which to examine genetic risk. Although PD risk may manifest in different cell types, here we focus on microglia. Microglia are unique in that they are both a glia and an immune cell. There is also an extensive body of literature demonstrating the involvement of microglia in PD through their role in prolonged CNS inflammation and the propagation of alpha-synuclein [9]. However, there is still a limited understanding of the mechanisms leading to microglia dysregulation in PD. Evaluating genetic risk mechanisms in microglia will elucidate how this cell type may contribute to the risk of developing PD.

The overall goal of this study was to find functional SNPs and match them to the genes that they affect, revealing processes that are altered in microglia during PD. We employed multiple strategies to prioritize 6,749 “SNPs of interest,” from the latest GWAS metanalysis [2], down to a subset relevant in microglia. We used ATAC-seq to identify SNPs located in regions of open chromatin in iPSC-derived microglia. We also used published ATAC-seq and H3K27ac ChIP-seq data sets from primary *ex vivo* microglia tissue [10]. These data sets were combined to identify SNPs located in consensus regions of open chromatin surrounded by H3K27ac marks, which demarcate active enhancers and promoters. Using this strategy, we report on a list of 73 SNPs as candidates for more in-depth mechanistic evaluation in microglia. We also found that multiple SNPs are located in the promoters of genes involved in the transport of substances in and between cells. Many of these genes are linked to lysosomal/autophagy functions, indicating that these processes may be impaired in microglia during PD.

## Materials and Methods

### iPSC-derived microglia cell cultures

Induced pluripotent stem cells (iPSCs) were obtained from ATCC (ACS-1019, DYS0100). For maintenance, they were cultured in StemFlex medium (ThermoFisher, A3349401) on Geltrex LDEV-free reduced growth factor basement membrane (ThermoFisher, A1413201). When cells reached 80% confluency, they were passaged using ReLeSR (STEMCELL technologies, 05872). For a detailed protocol of iPSC differentiation to microglia, see McQuade et al. [11]. Briefly, iPSCs were differentiated into hematopoietic progenitor cells (HPCs) over the course of 11 days using the STEMdiff Hematopoietic kit (STEMCELL technologies, 05310). Non-adherent cells were then collected and analyzed, using FLOW, for the presence of CD43. About 96% of these cells were CD43+. Roughly 100,000 HPCs per well were seeded into a 6-well plate in custom made media containing DMEM/F12 (ThermoFisher, 11320033), 2X insulin transferrin selenite (Gibco, 41400045), 2X B27 (Gibco, 17504044), 0.5X N2 (ThermoFisher, 17502048), 1X glutamax (Gibco, 35050061), 1X non-essential amino acids (Gibco, 11140050), 5 μg/mL human insulin (Sigma, 12643), 400 μM monothioglycerol (Sigma, M1753), and 1X pen strep (Gibco, 15140-122). For the first 24 days, new medium was added every other day starting on day 2, and supplemented with 100 ng/mL IL-34 (Peprotech, 200-34), 50 ng/mL TGFβ1 (Peprotech, 100-21), and 25 ng/mL M-CSF (Peprotech, 300-25), just before adding the media to cells. On day 25, all media was changed and supplemented with 100 ng/mL CD200 (Novoprotein, C311) and 100 ng/mL CX3CL1 (300-31), in addition to the three cytokines listed above. Fresh medium containing all five cytokines was added to the cells on day 27. On day 28, cells were cryopreserved in BamBanker (Wako, NC9582225). Expression of microglia specific markers Iba1 and TMEM119 were confirmed using immunofluorescence.

### ATAC-seq

Microglia were thawed and cultured for at least one week prior to an ATAC-seq experiment. Samples that yielded the best fragmentation started from a total of 10K, 31K, and 100K cells. The pre-specified number of cells were aliquoted into 1.5 mL tubes and centrifuged at 400 xg for 7 minutes. The supernatant was removed, and the cells were washed once with 50 μl ice-cold PBS. The cells were then resuspended in ice-cold Lysis Buffer containing resuspension buffer (1M Tris-HCl (final conc. = 10mM), 5 M NaCl (final conc. = 10 mM), 1M MgCl_2_ (final conc. = 3 mM), and nuclease-free H_2_O), 10% NP-40 (final conc. = 0.1% v/v), 10% Tween-20 (final conc. = 0.1% v/v), and 1% Digitonin (Promega, G9441) (final conc. = 0.01% v/v). Cells were then incubated on ice for 3 minutes. One mL of wash buffer (990 μl resuspension buffer + 10 μl Tween-20 (final conc. = 0.01% v/v) was added to each tube. The tubes were then inverted 3X gently and centrifuged at 500 xg for 5 minutes. For each sample, 10 μl of transposition mix (7.5 μl 2X TD Buffer (Illumina, FC-121-1030), 2.05 μl 1X PBS, 0.15 μl 10% Tween-20 (final conc. = 0.1 v/v), 1% Digitonin (final conc. = 0.01% v/v), and 0.15 nuclease-free H_2_O) was added. Five μl of ATM (Illumina, FC-121-1030) was then added separately to each sample. The samples were incubated for 60 minutes on a thermomixer at 1,000 rpm. Following incubation, the samples were placed on ice, and 5 μl of NT buffer (Illumina, FC-121-1030) was added to each tube to neutralize the tagmentation reaction. Tubes were then centrifuged at 300 xg at 20°C for 1 minute and incubated at room temp for 5 minutes. DNA purification was done using the Zymo clean and concentrator kit (Zymo, D4014).

For library generation, 5 μl of Illumina Nextera DNA unique Dual Indexes (Illumina, 20027214) plus 25 μl NEBNext High-Fidelity 2X PCR Master Mix (NEB, M0541L) was added to 20 μl of purified transposed DNA. The transposed fragments were amplified starting at 72°C for 5 minutes, 98°C for 30 seconds and then five cycles of 98°C for 10 seconds, 63°C for 30 seconds, and 72°C for 1 minute. qPCR was used to determine how many additional cycles to run on each sample. The PCR mix was composed of 5 μl of the partially amplified library from the previous step, 0.5 μl Illumina primer 1 (25 μM, 5’-AATGATACGGCGACCACCGA-3’), 0.5 μl Illumina primer 2 (25 μM, 5’-CAAGCAGAAGACGGCATACGA-3’), 0.75 μl 20X Eva Green, and 5 μl NEBNext High-Fidelity 2X PCR Master Mix. Cycle conditions were set to 98°C for 30 seconds, and 20 cycles of 98°C for 10 seconds, 63°C for 30 seconds, and 72°C for 1 minute. The R vs. cycle number was plotted on a linear scale. Additional cycles were calculated by determining the number of cycles needed to reach 1/3 of the maximum R. PCR was continued on the remaining partially amplified libraries for the appropriate number of cycles calculated in the previous step.

### Size selection and Sequencing of ATAC-seq Libraries

Library quantification, size selection, and sequencing were carried out by the genomics core at VAI. PCR amplified libraries were size selected for fragments 200-800 bp in length using double-sided SPRI selection (0.5x followed by 1x) with KAPA Pure beads (Kapa Biosystems). The quality and quantity of the finished libraries were assessed using a combination of Agilent DNA High Sensitivity chip (Agilent Technologies, Inc.), and QuantiFluor^®^ dsDNA System (Promega Corp., Madison, WI, USA). 75 bp, paired-end sequencing was performed on an Illumina NextSeq 500 sequencer using a 150 bp sequencing kit (v2) (Illumina Inc., San Diego, CA, USA) to produce a minimum of 50M paired-reads per library. Base-calling was done by Illumina NextSeq Control Software (NCS) v2.0, and the output of NCS was demultiplexed and converted to FastQ format with Illumina Bcl2fastq v1.9.0.

### Identification of ATAC-seq peaks

Four replicates of one iPSC-derived microglia cell line and thirteen replicates of ATAC-seq data from different primary microglia samples (published data) were used to find consensus ATAC-seq peaks. ATAC-seq peak data from primary microglia were obtained from dbGAP deposited by the Glass lab [10]. All data from iPSC-derived and primary microglia were processed the same way. Sequencing depth for iPSC-derived microglia was about 40-50 million, and read length was about 75 base pairs. For primary microglia, samples were sequenced to a depth ranging from 20-50 million reads, and the read length ranged from 47-76 base pairs [10]. Using Trimgalore, reads were trimmed or removed if they were below 20 bp in length. Forward and reverse reads for iPSC-derived microglia, and single-end reads for primary microglia were then aligned to the hg19 genome using the default settings for BWA v0.7.17 [12]. Multiqc v1.0 [13] was then run on all samples following alignment. Samblaster v0.1.24 [14] was used to sort and mark duplicate reads in bam files, and Samtools v1.9 [15] was used to remove duplicate reads and index the bam files. Peaks were then called using MACS2 v2.1.1 [16] default parameters. GenomicRanges v3.11 [17] was used to generate a consensus peak set (starting from narrowPeak files). We used peaks present in 3/4 samples for iPSC-derived microglia and peaks that were present in 10/13 primary microglia samples.

### PD risk SNPs and their overlap with ATAC-seq peaks

The list of 6,749 SNPs was obtained from the Nalls et al. bioRxiv version [18] from a supplemental file labeled “SNPs of interest tagging genes for functional inferences and networks analysis.” Using Bedtools, we searched for overlaps between the location of PD risk SNPs and ATAC-seq peaks from iPSC-derived microglia. The intersecting regions were then evaluated in IGV for SNPs that overlapped or were in close proximity (within 100 bp) of ATAC-seq and H3K37ac ChIP-seq peaks from primary microglia. In this analysis, we found a total of 73 SNPs, which we then ranked by GWAS p-value (Supplemental Table S1).

### Analysis for correlation of degree from allelic balance vs. effect size

To calculate the degree from allelic balance for each SNP, we took the absolute value of 0.5 minus the allele frequency. To calculate the effect size of each SNP, we took the absolute value of 1 minus the odds ratio. The association between the degree from allelic balance and the log(odds) was estimated using R v3.6.0 (https://cran.r-project.org/) via a robust linear regression and MM estimation [19].

### SNP annotation and network analysis

The locations of the 73 SNPs in ATAC-seq peaks were annotated using ChIPseeker v3.11 [20]. We then took the top 38 genes from supplemental Table 1, with SNPs in their promoters, and entered them into the MSigDB database [21, 22] to evaluate any overlap with GO or immunologic gene sets. Default parameters, including an FDR q-value less than 0.05, were used. The GO gene sets that were manually selected are derived from the Gene Ontology Resource [23, 24]. The gene sets for the immunologic categories come from individual studies [25-28]. Their GSE numbers can be found in Figure 3 B.

## Results

### ATAC-seq peaks in iPSC-derived microglia vs. primary microglia

We evaluated genetic risk in microglia using the iPSC-derived microglia model developed by McQuade et al. [11]. Following the differentiation of iPSCs to mature microglia, we performed ATAC-seq to map genome-wide regions of accessible chromatin. iPSC-derived model systems represent a promising alternative to primary tissue due to their relative ease of creation and experimental manipulation. However, culturing cells in an *in vitro* environment may affect the chromatin landscape [10]. Therefore, to augment our newly generated ATAC-seq data from iPSC-derived microglia, we also obtained ATAC-seq data from 13 *ex vivo* primary microglia samples published by the Glass laboratory. [10]. For these samples, microglia were isolated from the brain tissue of 13 different patients ranging in age from 2 to 17 years. These samples were also obtained from various regions, including the temporal cortex, the frontal cortex, the occipital cortex, and the cerebellum. We combined the data sets to identify active chromatin regions in iPSC-derived microglia shared with a heterogeneous population of primary microglia. Presumably, these common loci are relevant to the function of a broad range of microglia types. Peak sets for iPSC-derived microglia (both overlapping and non-overlapping with primary microglia) can be found in the supplement (Supplemental Table S2 and S3).

Many ATAC peaks are shared between iPSC-derived microglia and primary microglia. Out of 73,276 peaks present in iPSC-derived microglia, 60,139 (~82%) overlap with primary microglia (Figure 1 A). An example of peak consistency at a particular locus is shown in Figure 1 B around *CX3CR1*, a gene known to be expressed in microglia. For subsequent data analyses, we used the peaks present in both iPSC-derived and primary microglia.

**Figure 1:**
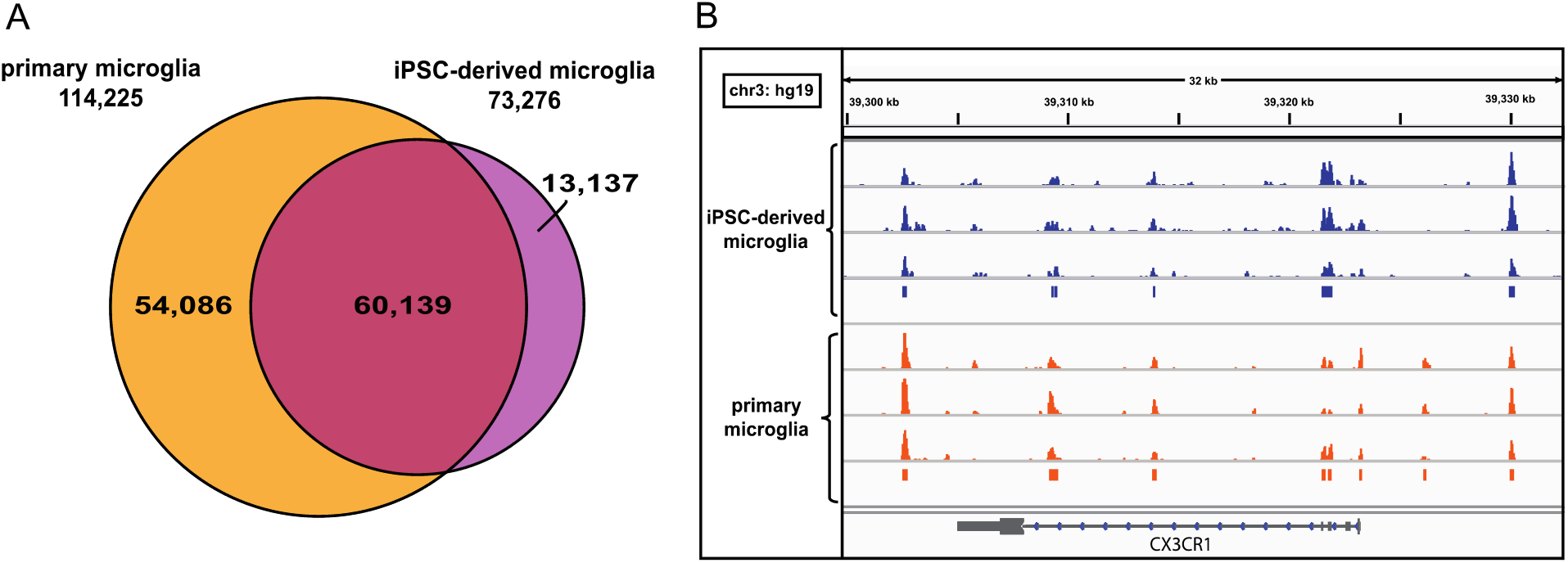
Comparison of ATAC-seq peaks in iPSC-derived microglia to primary microglia. (A) Euler diagram of the overlap between primary and iPSC-derived microglia ATAC-seq peaks. (B) IGV screenshot of 3 replicates of one iPSC-derived microglia cell line (blue) and three different primary microglia cell lines (one replicate of each) (orange) at the CX3CR1 locus. Blue and Orange bars represent peaks called by MACS2 [16].

### Finding candidate functional SNPs

We identified SNPs that are potentially functional by searching in regions of open chromatin (mapped using ATAC-seq). In addition to using ATAC-seq, we also obtained published H3K27ac ChIP-seq data [10] from 3 primary microglia samples to further identify SNPs within regulatory regions. Although SNPs likely function in multiple ways, those that reside in these regions may function by disrupting transcription factor binding, leading to changes in gene expression [29]. To identify such SNPs, we first overlaid the location of PD risk “SNPs of interest” from Nalls et al. [18] with iPSC-derived and primary microglia ATAC-seq peaks. Out of 6,749 SNPs, 73 were located in ATAC peaks and were in or within 100 bp of an H3K27ac ChIP-seq peak present in primary microglia (Supplemental Table S1). We observed a significant positive correlation (p-value = 3.24 x 10-3) between the degree of deviation from allelic balance and effect size on risk (Supplemental Figure S1), indicating, as expected, that rarer SNPs have higher penetrance and a more substantial influence on PD risk.

From the 73 SNPs in ATAC peaks, we highlight six top candidate risk SNPs (Figure 2) based on GWAS p-value and magnitude of effect on risk (odds ratio). The top-ranking SNP based on both p-value (2.208×10^−27^) and odds ratio (1.97) is rs12726330. It is located within the promoter of *SLC50A1*, which is a gene that encodes for the protein SWEET1 [30], whose function is not completely known [31]. The second candidate rs2737004 (p-value = 7.60×10^−11^), is located within a potential enhancer in an intron of *SNCA. SNCA* codes for the protein alpha-synuclein, which is mostly known as a presynaptic neuronal protein that regulates neurotransmission [32]. Rs144814361 is located within the 5’UTR/promoter of the *BAG3*. This SNP has the third-lowest p-value (9.07×10^−11^) and the second-highest odds ratio (1.55). *BAG3* is a gene that is critical for preventing apoptosis during aging and under conditions of stress [33]. Rs3813020 has the fourth-lowest p-value (2.05×10^−10^). It is in the promoter of *FBXL19*. FBXL19 is a member of the Skp1-Cullin-F-box family of E3 ubiquitin ligases that regulates the ubiquitination and degradation of inflammatory cytokines like TNF‐α, IL‐1β, and IL‐6 [34]. Rs4889599 (p-value = 7.34×10^−10^) is located in the promoter of *SETD1A*. SETD1A is a histone methyltransferase complex component that regulates mono, di, and trimethylation of histone H3 at Lys4 [35]. Lastly, rs823114, with the 7^th^ lowest p-value (4.35×10^−09^), and a less significant SNP, rs7536483, are located in the promoter of *NUCKS1* (Nuclear casein kinase and cyclin-dependent kinase substrate 1). NUCKS1 is a chromatin-associated protein with a role in the DNA damage response, DNA repair, metabolism, and inflammatory and immune responses, but its overall function is still unknown [36]. Other than rs4889599, we observed that these SNPs are located in ATAC-peaks that are flanked by H3K27ac ChIP-seq peaks. This pattern is what we typically consider optimal when evaluating candidate loci because it is indicative of nucleosome displacement, likely due to transcription factor occupancy where the ATAC-seq peak is in the middle of the H3K27ac peak.

**Figure 2:**
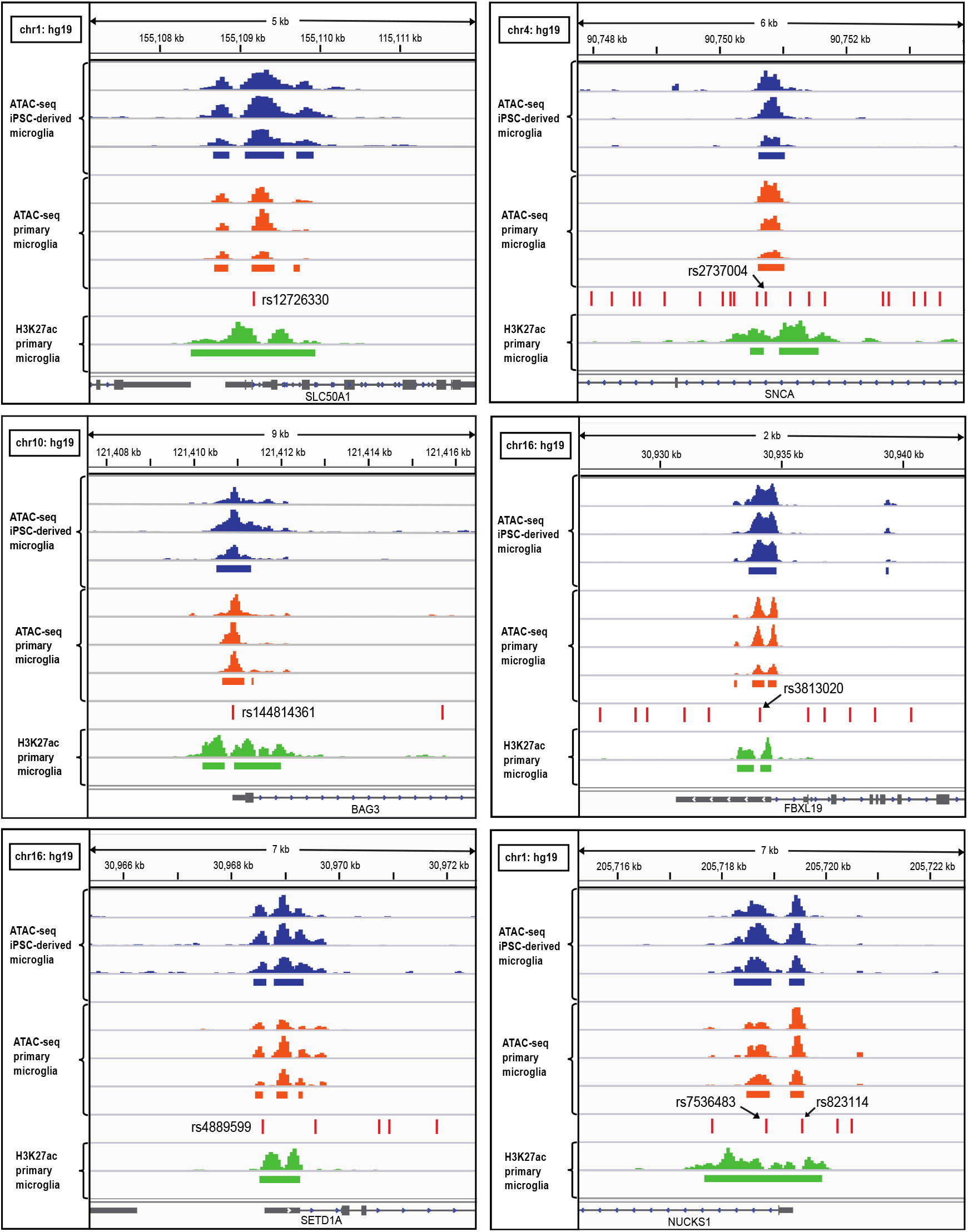
Six candidate risk SNPs. IGV screenshot of top candidate risk SNPs (red lines marked with arrows). These SNPs overlap peaks from iPSC-derived microglia (blue) and primary microglia (orange). They are also surrounded by H3K27ac histone marks (green).

**Figure 3:**
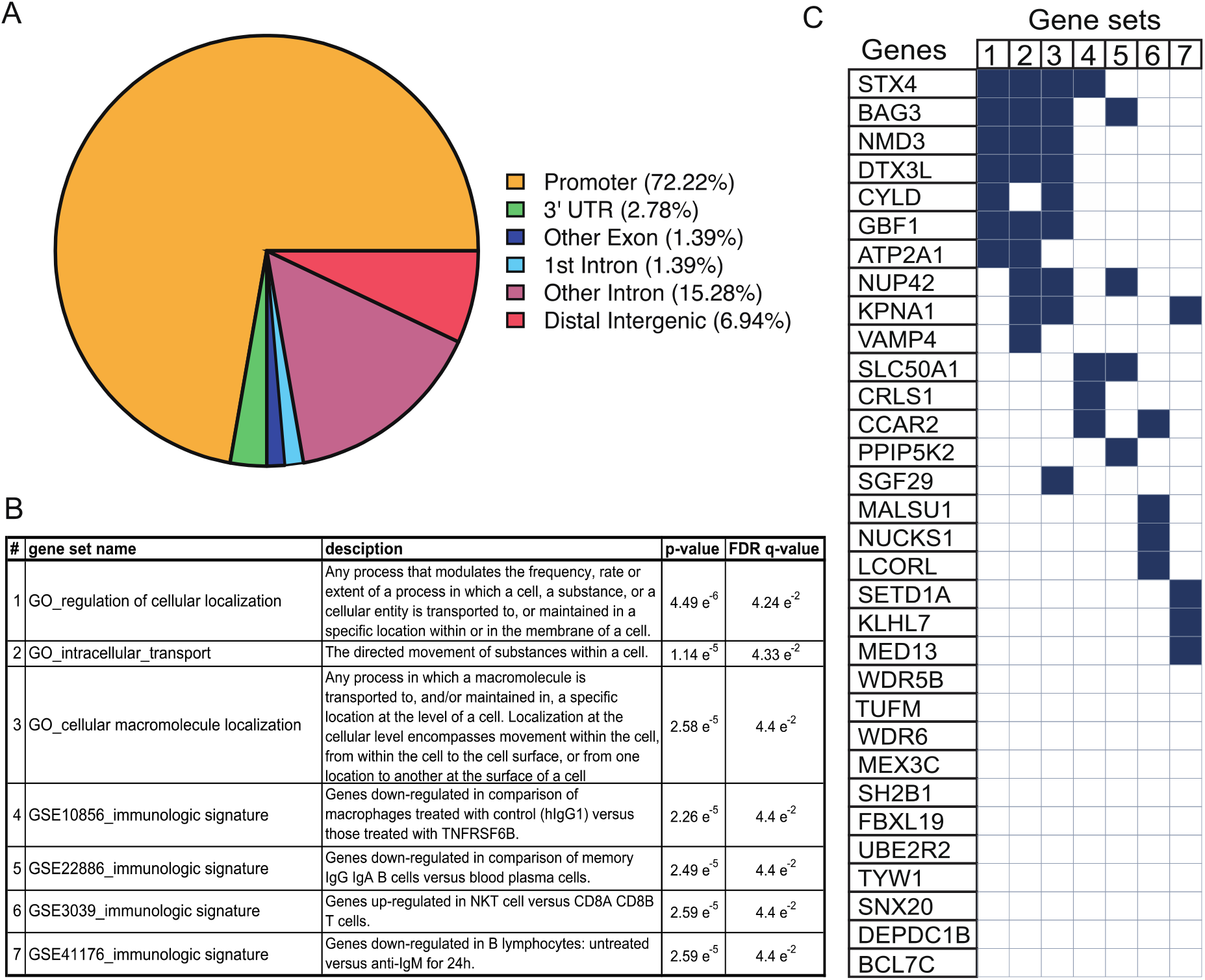
Assessment of SNPs’ location in ATAC peaks and network analyses of genes containing SNPs in their promoters. (A) Pie chart showing the location of 73 risk SNPs in consensus primary and iPSC-derived microglia ATAC peaks. The majority of SNPs (53) are located in promoters. (B) Table, from MSigDB [22, 44], of the categories/gene sets that significantly overlap genes with SNPs in their promoter. (C) Diagram showing which genes overlap which categories from the MSigDB analysis. In part B, the numbers in the first column of the table correspond to the gene set number in the figure in part C.

We used MotifbreakR to evaluate which SNPs potentially regulate gene expression via altered transcription factor binding (Supplemental Figure S4) [37]. Figure 3 shows the top three transcription factors, based on the strongest allele-specific effect on binding, for the top four ranking SNPs in Supplemental Table S1. Allele-specific binding of these transcription factors may affect the expression of SLC501A, SNCA, BAG3, FBXL19, or multiple genes in cis or trans. Some transcription factors like RAD21, ZNF143, and CTCF regulate gene expression through the facilitation of chromatin looping [38-40]. Other transcription factors in Figure 3, like ETS1, IRF3, and REST, have diverse functions, including developmental regulation, regulation of genes involved in cellular responses and immunity, and epigenetic remodeling, respectively [41-43].

### Multi-SNP network analysis of SNPs in promoters

As a first step to identify gene networks that may be altered in microglia during PD, we mapped the location of the 73 “SNPs of interest” in ATAC-seq peaks. Unexpectedly, the majority, 53 (~72%), were located in promoters (Figure 3 A). Although each SNP may affect multiple genes, the most likely risk gene for this set of SNPs is the gene linked to those promoters. Thus, we took the top ranking 38 genes with SNPs in their promoters (listed in Supplemental Table S1) and computed overlaps with gene sets from MSigDb [21, 22, 44]. Gene sets queried for overlap originate from the Gene Ontology (GO) Network and from individual studies that have curated immunologic signatures categories (see methods for more details). Before running the analyses, a significance threshold of FDR q-value < 0.05 was set. Gene sets belonging to immunologic signatures categories showed a significant overlap. The most significant categories were related to GO gene sets involved in cellular localization of substances/macromolecules. Many of the proteins in these categories, like STX4, BAG3, CYLD, VAMP4, and others, have known roles in autophagy and lysosomal protein degradation [45-49].

## Discussion

Here we aimed to find a subset of PD-associated SNPs located in regions of active regulatory DNA in microglia in order to identify functional risk SNPs in this cell type. In doing so, we substantially narrowed down 6,749 PD associated SNPs to a more tractable list of 73 SNPs. All of these SNPs are predicted to disrupt transcription factor binding motifs, suggesting that they may be functional via allele-dependent gene expression that consequently increases risk for PD.

Although a comprehensive analysis of the 73 SNPs in ATAC-seq peaks is our longterm goal, we have highlighted some top-ranking SNPs as candidates for immediate mechanistic follow up. For brevity, we only discuss two SNPs within genes that have been previously linked to PD. Rs2737004 is located within *SNCA* in a likely intergenic enhancer and may therefore affect the expression of the protein (alpha-synuclein) encoded by *SNCA*. Expression data (eQTL data) from GTEx does indeed show that the GG genotype of rs2737004, with G being the allele that carries the risk, is associated with increased expression of *SNCA* in the cortex. However, these findings have yet to be validated in microglia. In terms of a mechanism that may explain the allele-specific expression of *SNCA*, rs2737004 disrupts three separate binding motifs for CTCF (Figure 4 and Supplemental Table S4). CTCF can serve as an activator, a repressor, or an insulator, to block the interaction between enhancers and promoters [50]. Alterations in CTCF binding may, therefore, allow for the unregulated expression of *SNCA* or other genes.

**Figure 4:**
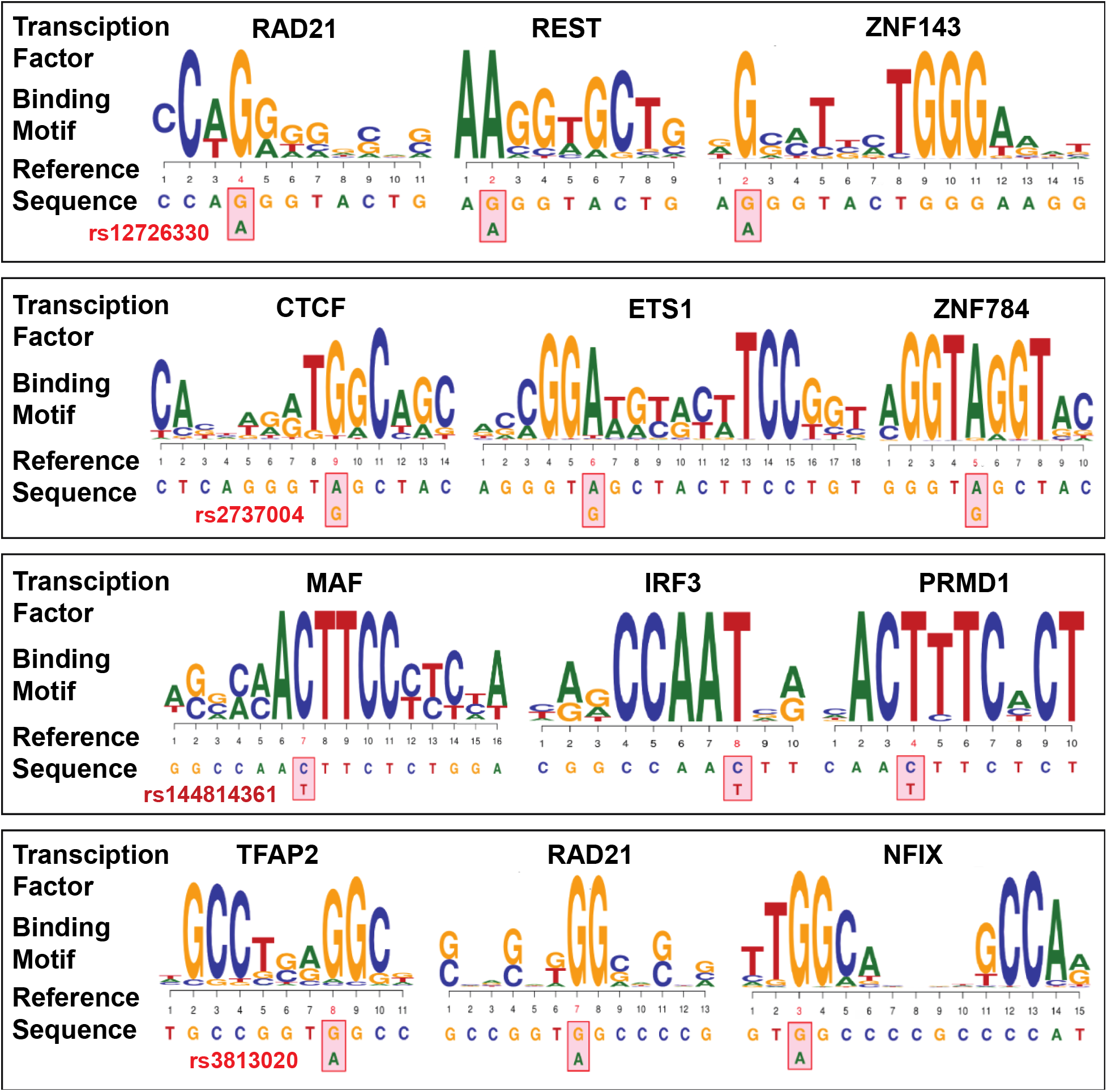
Transcription factors disrupted by risk SNPs. For each of the four top SNPs in Supplemental Table S1, three consensus DNA binding motifs are shown. These motifs are predicted to have allele-specific protein binding affinity.

Alterations in the expression level and the sequence of alpha-synuclein have been linked to familial and sporadic forms of PD [51, 52]. However, many aspects of alpha-synuclein’s functions are still under investigation. Alpha-synuclein is known to function within microglia, though its roles in neurons are usually emphasized in the literature. Studies have shown that alpha-synuclein can act as a chaperone through its interaction with Hsp70 [53, 54]. Hsp70 plays an essential role in protein quality control under conditions of stress [55]. It is therefore likely to be an important protein in microglia as this cell type is highly responsive to any changes that disturb homeostasis in the brain [56]. Indeed, Hsp70 can enhance phagocytosis by macrophages [57]. In addition, impairment of HSP70/HSP90 pathways leads to increased protein aggregation [58]. Thus, although not yet shown, alterations in the expression of *SNCA* may indirectly affect the Hsp70-mediated process of protein clearance in microglia.

Another strong candidate for follow up studies is rs144814361, located in the promoter of *BAG3*. BAG3 (Bcl associate athanogene 3) is a member of the BAG family of antiapoptotic proteins that, like alpha-synuclein, functions as a co-chaperone to regulate the activity of Hsp70 [33]. One of its key functions is to coordinate the selective degradation of misfolded and aggregated protein through selective macro-autophagy [33, 59]. This process is especially critical as an adaptive response under conditions of stress or aging [46]. BAG3 is linked to multiple age-related neurodegenerative disorders, including ALS, Alzheimer’s disease, and Parkinson’s disease [33, 60], and plays a protective role by removing disease-associated protein aggregates [60-62].

Work described by Cao et al. showed that *BAG3* expression was increased in the midbrain of *SNCA*_A53T_ transgenic mice and in cultured neuronal cells overexpressing wild-type *SNCA* [63]. The authors also found that *BAG3* overexpression enhanced autophagy and the clearance of alpha-synuclein. This study demonstrates the relevance of BAG3 as a vital mediator of protein quality control during stress and aging in neurons. However, BAG3’s involvement in autophagy and lysosomal protein clearance in microglia has yet to be determined.

The role of both SNCA and BAG3 as co-chaperones to Hsp70 suggests that stress-related protein clearance may be a mechanism in microglia that is influenced by multiple genetic variants. Further experimental validation is needed to determine if rs2737004 and rs144814361 affect the expression of *SNCA* and *BAG3*, respectively, and if so, what the observable effects are on microglia function.

For our network analysis, we searched for genes, with risk SNPs in ATAC peaks at their promoters, that overlapped immune pathways, due to the importance of microglia in the CNS immune response. A number of the genes singled out by this analysis are interesting due to their relevance to inflammatory cells. We only discuss *SLC50A1*, which has the most significant p-value out of the 73 SNPs in Supplemental Table S1. *SLC50A1* encodes for the sugar transporter SWEET1. There is little known about its function [31], but as a glucose transporter, it could significantly impact the function of immune cells such as microglia. Activation of immune cells requires increased glucose utilization, which is in part mediated by the upregulation of glucose transporters [64]. Alterations in *SLC50A1* expression may consequently affect microglia activation, but this has yet to be validated.

Consistent with the role of SNCA and BAG3 in protein clearance, multiple other genes from our network analysis (Figure 3) are linked to lysosomal and autophagy functions. For example, STX4 facilitates the fusion of autophagosomes to the plasma membrane [49]. CYLD, a deubiquitinase specific for lysine63-linked polyubiquitin chains, has been shown to play a role in neurodegeneration by removing ubiquitin from synaptic proteins, preventing their autophagic removal [65]. NMD3 is a 60S ribosomal subunit protein that, when inhibited, may induce autophagy and, when hyperactive, may decrease autophagy [66, 67]. Loss of VAMP4, a vesicle-associated membrane protein, has been linked to the accumulation of dysfunctional lysosomes and reduced autophagic lysosomal degradation [48]. In addition to the genes mentioned above, other genes from this network analysis are associated with autophagic and lysosomal processes. Most of the SNPs in these genes’ promoters have eQTL values that show expression levels of these genes in various tissues is allele-dependent, implying that autophagy and lysosomal pathways may be affected by many of these SNPs.

Prior studies indicate that autophagy/lysosomal processes are altered in PD, and they point to specific genes such as *PINK1, LRRK2, GBA*, and *ATP13A2* that may be involved, but much of these findings are from multiple cell types or neuronal populations [68]. For microglia, our data shows that, except for a SNP in *LRRK2*, there are no “SNPs of interest” located in ATAC peaks at regulatory DNA at well-known PD associated lysosomal genes. Instead, the genes listed in Figure 3 may be the ones specifically affected in microglia and are the ones involved in autophagy and lysosomal dysfunction in this cell type during PD. Indeed, microglia have been shown to play an important role, more so than other cell types, in the degradation of alpha-synuclein, which is mediated through their lysosomal and autophagy functions [69]. However, there are still a limited number of studies investigating the role of lysosomal and autophagic dysfunction in microglia during PD. Further studies are needed to confirm if the genes in Figure 3 or other genes belonging to the same network are affected by the risk variants we highlight in this study. This additional work will help to shed light on the contribution of microglia to PD risk.

There is much work to be done to validate the target risk genes of the 73 SNPs identified in this study. When defining risk genes, as we have done here, a common approach is to take the nearest genes to the SNP. Whereas this approach has provided substantial insight into PD related pathways, it may not reveal the full extent of gene targets because a SNP may affect genes multiple kilobases away or even on different chromosomes [70]. Therefore, it will be essential to validate our preliminary findings with further experimental follow-up studies. Techniques such as genomic editing, using CRISPR/Cas9 to either create a small lesion at the SNP or exchange one allele for the other, followed by RNA-seq, allows for the evaluation of allele-specific effects on gene expression. High-resolution chromatin conformation capture assays like 3C and 4C can then be utilized to confirm regulatory DNA-gene contacts affected by a risk SNP. The identified risk genes can then be evaluated, as a set, for any associations with biological processes, revealing potential mechanisms for further hypothesis testing.

Our data provide a starting point for dissecting genetic risk in microglia. Although multiple gene networks, such as those involved in inflammation, may be affected by PD risk SNPs, our assessment of individual SNPs like the ones at *BAG3* and *SNCA*, and our multi-SNPs network analysis, supports the hypothesis that autophagy/lysosomal processes may be altered within microglia in PD. The extent to which microglia contributes to defects in autophagy and lysosomal functions in PD remains an opened question that should be addressed due to potential impacts on alpha-synuclein aggregation. We advocate for more post-GWAS testing of these risk variants to uncover genetic factors that increase PD risk. There are currently no treatments to modify the progression of PD. Additional studies that build on our findings will help understand the complex genetic etiology of PD and identify alternative disease-modifying targets.

## Acknowledgments

Van Andel Institute provided the facilities and general support to conduct the study. Marie Adams and the rest of the Genomics Core conducted and advised on nextgeneration sequencing. Rachael Sheridan and the Flow cytometry core conducted the Flow analysis. Lastly, Zach Madaj provided support for statistical methods and analysis.

## Author Contributions

### A Booms

Did all experimental work and wrote the paper.

### SE Pierce

Contributed analyses and edited the manuscript.

### GA Coetzee

Directed the lab, instigated the project/ideas/analyses and edited the manuscript.

All authors read and concurred with the final manuscript.

## Supplemental Figure Legends

**Supplemental Table S1. 73 PD risk SNPs in ATAC-seq peaks in microglia.** The SNPs in this table are ranked based on significance (GWAS p-value for risk). SNPs are listed with GWAS statistics from Nalls et al. [2]. ChIPseeker annotations are listed in the last column.

**Supplemental Table S2. ATAC-seq peaks from iPSC-derived microglia**

**Supplemental Table S3. ATAC-seq peaks shared by iPSC-derived and primary microglia**

**Supplemental Table S4. MotifbreakR results for 73 SNPs.** Transcription factors that are not expressed in microglia were excluded.

---

**Supplemental Figure S1.**
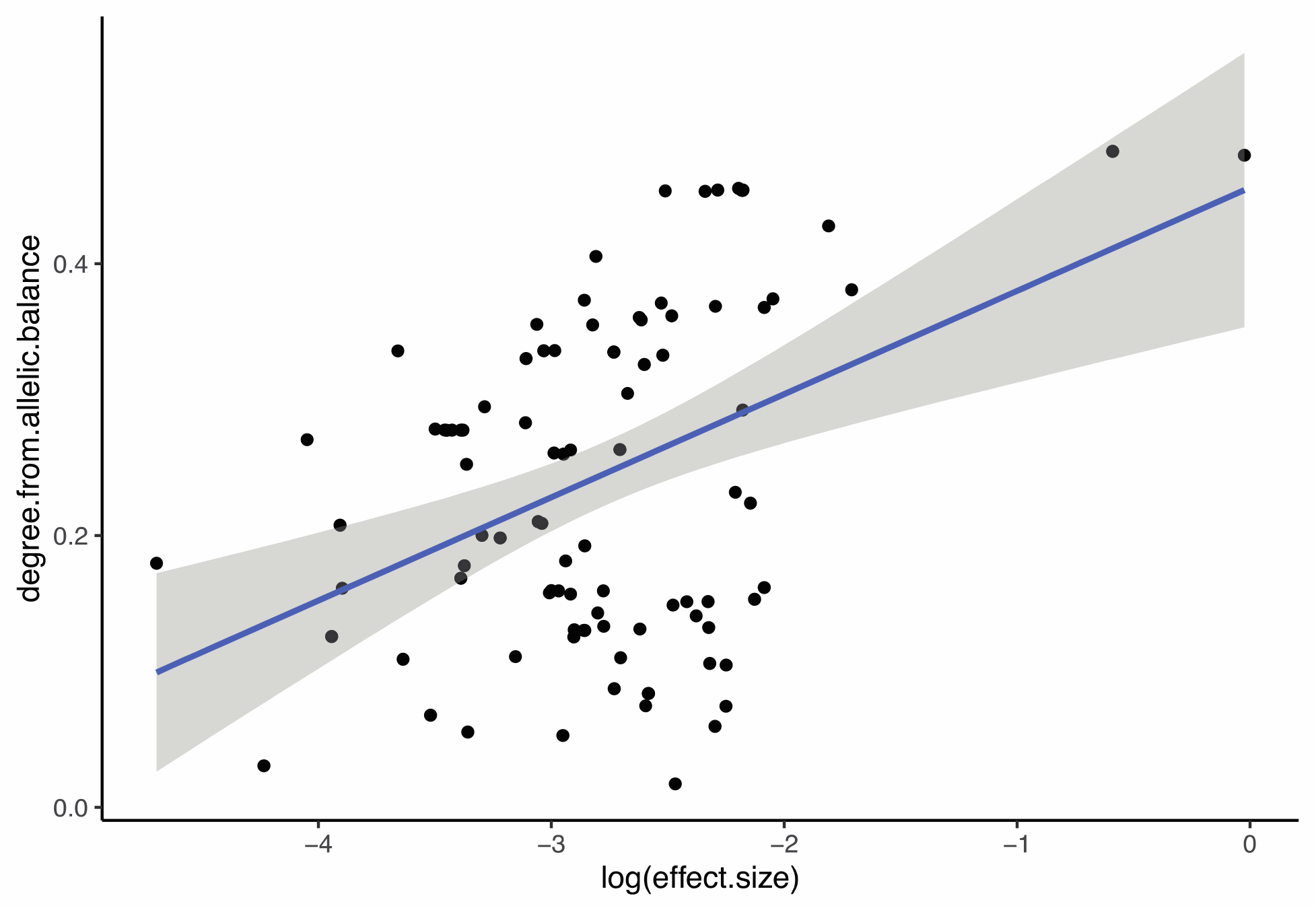
Relationship between the degree from allelic balance and effect size of 73 SNPs in ATAC-peaks. The absolute values of allele frequencies normalized to 0.5 were plotted against the absolute values of odds ratios normalized to 1.

## Notes

### Competing Interest Statement

The authors have declared no competing interest.

